# descSPIM: Affordable and Easy-to-Build Light-Sheet Microscopy for Tissue Clearing Technique Users

**DOI:** 10.1101/2023.05.02.539136

**Authors:** Kohei Otomo, Takaki Omura, Yuki Nozawa, Yuri Saito, Etsuo A. Susaki

## Abstract

Despite the easier use of multiple tissue clearing techniques in recent years, poor access to adequate light-sheet fluorescence microscopy remains a major obstacle for biomedical end users. Here, we propose a solution by developing descSPIM (*des*ktop-equipped SPIM for *c*leared specimens) as a low-cost ($20,000–50,000), low-expertise (one-day installation by a non-expert), yet practically substantial do-it-yourself light-sheet microscopy. Academically open-sourced (https://github.com/dbsb-juntendo/descSPIM), descSPIM allows routine three-dimensional imaging of cleared samples in minutes.

---

Since the early attempts(*1*), tissue clearing has become the gold standard for volumetric tissue, organ, and body imaging. The maturation has supported numerous scientific discoveries(*2, 3*). Highly efficient tissue clearing techniques such as DISCO(*4*), CUBIC(*5, 6*), and SHIELD(*7*), which enable 3D imaging of organs and the entire body using single-photon light-sheet fluorescence microscopy (LSFM), have provided detailed protocols or have already been commercialized, enabling end-users to easily approach these techniques.

However, the limited accessibility of LSFM remains a major obstacle to the widespread utilization of tissue clearing techniques (**Fig. 1a**). Although several systems are commercially available and provide a highly usable configuration and interface, they are prohibitively costly (>$500k). Also, biomedical end-users cannot easily access cutting-edge custom-built microscopy which primarily aim to achieve technical leadership. mesoSPIM(*8*) was designed as a more user-oriented system, reducing the barrier. Nevertheless, its installation and maintenance still require a degree of expertise in optics and microscopy construction (**Fig. 1a**; **Supplementary Fig. 1**). Other open-source projects in the early days, such as openSPIM(*9*) and OpenSpin microscopy(*10*) were more optimized for live imaging of small animals but not suitable for specimens treated with modern tissue clearing methods (**Supplementary Fig. 1**). Here, we present *descSPIM* (descSPIM-basic) as a simple, low-cost, yet practically effective system particularly designed for tissue clearing technique users in biomedical fields who are unfamiliar with microscopy systems.

**Fig. 1.**
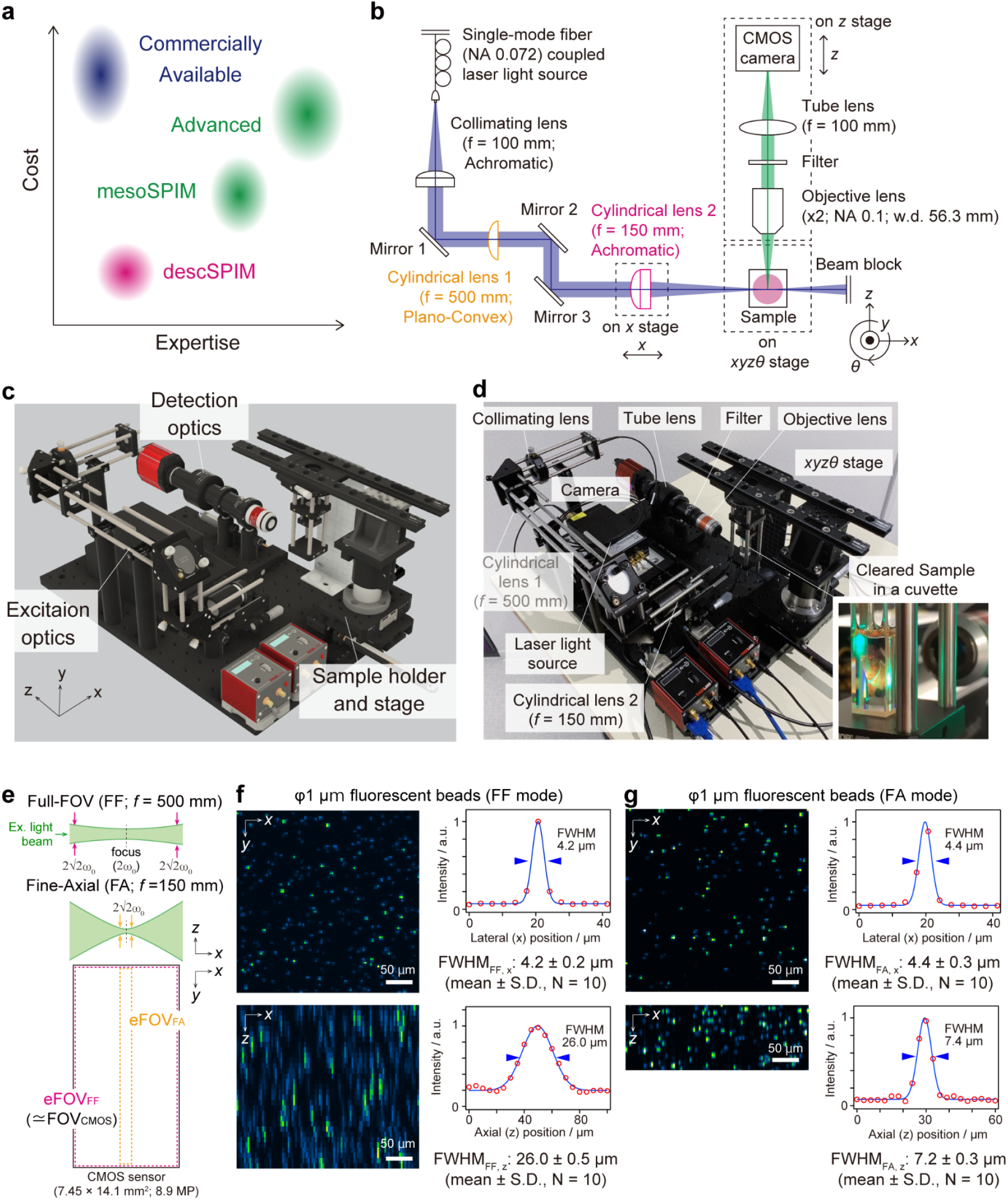
descSPIM concept, components, and optical specifications. **a**. The target of descSPIM on the expertise and cost axes, where the simultaneous achievement of thorough simplification and practical quality is required. **b**. descSPIM system overview. A one-sided light-sheet illumination formed by a collimated lens and a cylindrical lens comes from the right side of the sample. Either a cylindrical lens (f = 500 mm or f = 150 mm) is used for each illumination mode (FF and FA), respectively. A manual linear translation stage is attached to the latter cylindrical lens for the TLS imaging. The detection path is composed of a 2X objective lens, tube lens, and CMOS camera, with an effective integration magnification of 1X (3.45 μm × 3.45 μm of the pixel size). **c**. Schematic overview of the descSPIM-basic. **d**. Overview of the actual descSPIM-basic instrument. (inset) The cleared sample can be placed in a four-sided transmission cuvette and directly illuminated for imaging. No oil chamber was adopted. **e**. Two illumination modes adopted to the descSPIM-basic by switching the cylindrical lenses. FF covers the whole CMOS sensor area (7.45 mm in width). FA provides finer axial resolution while covering ∽11% (830 μm of the Rayleigh length in width) of the sensor area. eFOV: effective field of view. **f** and **g**. Estimation of lateral and axial resolutions by PSFs (evaluated with images 1 μm beads images). Comparable xy resolution was achieved by the two illumination modes. FA provides approximately 7 μm of axial FWHM, over three-times finer than FF mode.

The invention of descSPIM necessitated a fundamental redesign of all configurations for functionality and minimization. The system basically adopted the original SPIM configuration for illumination optics(*11*), forming a one-sided light-sheet illumination with minimal equipment (**Fig. 1b**). All the components for the excitation/detection optics and stages (approximately 90 parts) were aligned on a 300 mm x 450 mm breadboard (**Fig. 1c, d**). They can be purchased from a single vendor (Thorlabs) for the user’s convenience, which is in contrast to other open-source projects using several custom parts (**Supplementary Table 1**; **Supplementary Fig. 1**).

A single-mode fiber-guided laser light was collimated with an achromatic convex lens, then formed into sheet illumination by a cylindrical lens. Users can choose color variations according to their experimental purpose. We used a four-color laser light source (Cobolt Skyla 488/515/561/647 nm) for multichannel imaging.

The sample stages included manual *x, y* and *θ* stages to reduce costs, and a motorized *z* stage for imaging and focusing. The cleared sample was placed in a four-sided transmission cuvette so that the users can handle it without an imaging oil chamber (**Fig. 1c, d**). One of the z stages moved the sample, while the other shifted the detection optics. The latter stage was required to get into a focused position and to prevent defocusing during imaging, which was caused by the change in the distance ratio between the clearing reagent with a high refractive index (RI) and the air with a RI of 1.00 (**Supplementary Fig. 2**).

The detection optics included an objective lens with a low numerical aperture (NA) and long working distance (WD) (Thorlabs TL2X-SAP) combined with a tube lens and a CMOS camera with a rectangular sensor (size 14.131 mm × 7.452 mm). The sensor shape and size allowed the effective width of the light sheet to cover the whole sensor area as a field-of-view (FOV) (**Fig. 1e, Supplementary Fig. 3a**). The resulting images, acquired at an effective integration magnification of 1X, had a pixel size of 3.45 × 3.45 μm^2^.

For the practical use of the basic configuration, two types of cylindrical lenses were adopted to create wider or axially finer illumination (**Fig. 1e; Supplementary Fig. 3a**). Full-FOV (FF) illumination was generated by one of the cylindrical lenses (f = 500 mm) with an axial spatial resolution of 26 ± 0.5 μm (**Fig. 1f**) and an effective field of view (eFOV) (defined as 2 × Rayleigh length(*12*), **Supplementary Fig. 3b**) that largely covers the CMOS sensor area (**Supplementary Fig. 3c**). The other cylindrical lens with a shorter focal length (f = 150 mm) yielded fine-axial (FA) illumination with the axial spatial resolution of 7.2 ± 0.3 μm with a narrower eFOV (**Fig. 1g**; **Supplementary Fig. 3d**). As for lateral (*x*-y) resolution, comparable values were achieved by the two illumination modes (4.2 ± 0.2 μm for FF mode and 4.4 ± 0.3 μm for FA, respectively) (**Fig. 1f, g**). Here, the spatial resolution was estimated by full-width of half maximum (FWHM) values of point spread function (PSF) that was defined from the intensity profile of the signals from a Φ1 μm bead. In the tissue sample images in this paper, we considered practical z-step ranges to be roughly half of FWHM, with a minimum of 10 or 5 μm for FF and FA modes, respectively.

The FWHM values were also utilized to extrapolate the literature-defined effective NA value of cylindrical lenses for SPIM(*12*) (**Supplementary Fig. 3b**). We determined the effective NA values to be 0.007 for the FF mode and 0.033 for the FA mode. Based on the NA value, the eFOV in FA mode is approximated as 830 μm (**Supplementary Fig. 3b**), which is consistent with the eFOV estimation from an image of 1 μm beads (**Supplementary Fig. 3c**). Therefore, the cylindrical lens used in FA mode yields a sheet illumination near to the approximation. Since the ratio of NA values is equalized to the ratio of focal length, the effective NA of the cylindrical lens used in FF mode based on the NA of the cylindrical lens in FA mode (0.033) should be 0.01 (0.033 × 150 mm/500 mm), which is inconsistent with the effective NA of the cylindrical lens in FF mode (0.007). This discrepancy could be attributable to the aberration in the plano-convex lens used in FF mode, which led to a lower effective NA. Consequently, the cylindrical lens in FF mode generated a homogeneous light sheet illumination (approximately 26–29 μm of FWHMs) across the CMOS sensor area (**Supplementary Fig. 3b**).

Data acquisition and processing procedures are summarized in **Supplementary Fig. 4**. The basic system does not require users to deploy any custom programs for operations; it can be used only with the device-associated software. Two actuators should be moving synchronously at distinct velocities. The synchronous speed correction value (the relative velocity of two actuators) can be determined by the method in **Supplementary Fig. 4**. Inspired by the MOVIE method(*13*), a z-stack image was rapidly collected as time-lapse (*xy*-*t*) data by the continuous moving of the actuators, and then the data was converted into *xy*-*z* format. This method enabled minute-order volumetric imaging per image stack.

As representatives, we collected a CUBIC-cleared and propidium iodide (PI)-stained mouse hemisphere and a 2 mm-thick mouse brain slice acquired with FF and FA modes, respectively (**Fig. 2a-d**). The speed was 3-6 minutes/stack, depending on the illumination mode. To homogenize the illumination intensity in the *y* direction, we adopted a flat-field correction (FFC) of gaussian illumination(*14*) (**Supplementary Fig. 5**) with a parallelly obtained image of the dye solution (1 μM fluorescein in CUBIC-R). We further took up the tiling light-sheet method (TLS)(*15*) into FA mode imaging for gaining uniform axial resolution over the sample thickness (**Supplementary Fig. 6**). The resulting axial resolutions for the nuclei, estimated by FWHM of PSF, were consistent with the results by Φ1 μm beads (**Fig.1 f, g** and **Fig. 2b, d**). In the image obtained by FF mode, the axial FWHMs at the center and edge positions of the FOV were comparable (29.5 ± 0.2 μm and 27.0 ± 0.1 μm), again suggesting the homogeneous light-sheet thickness across the eFOV (**Fig. 2a, b**). As for the image with FA mode, the elongation of axial FWHMs compared with lateral FWHMs (11.4 ± 0.7 μm in axial and 8.2 ± 0.8 μm in lateral, approximately 1.37× elongation) was reduced compared with the axial elongation obtained with Φ1 μm beads (7.2 ± 0.3 μm in axial and 4.4 ± 0.3 μm in lateral, approximately 1.64× elongation) (**Fig. 1g** and **Fig. 2d**), indicating that the FA mode provides a quality of image close to the isotropic resolution for the nuclei-size objects.

**Fig. 2.**
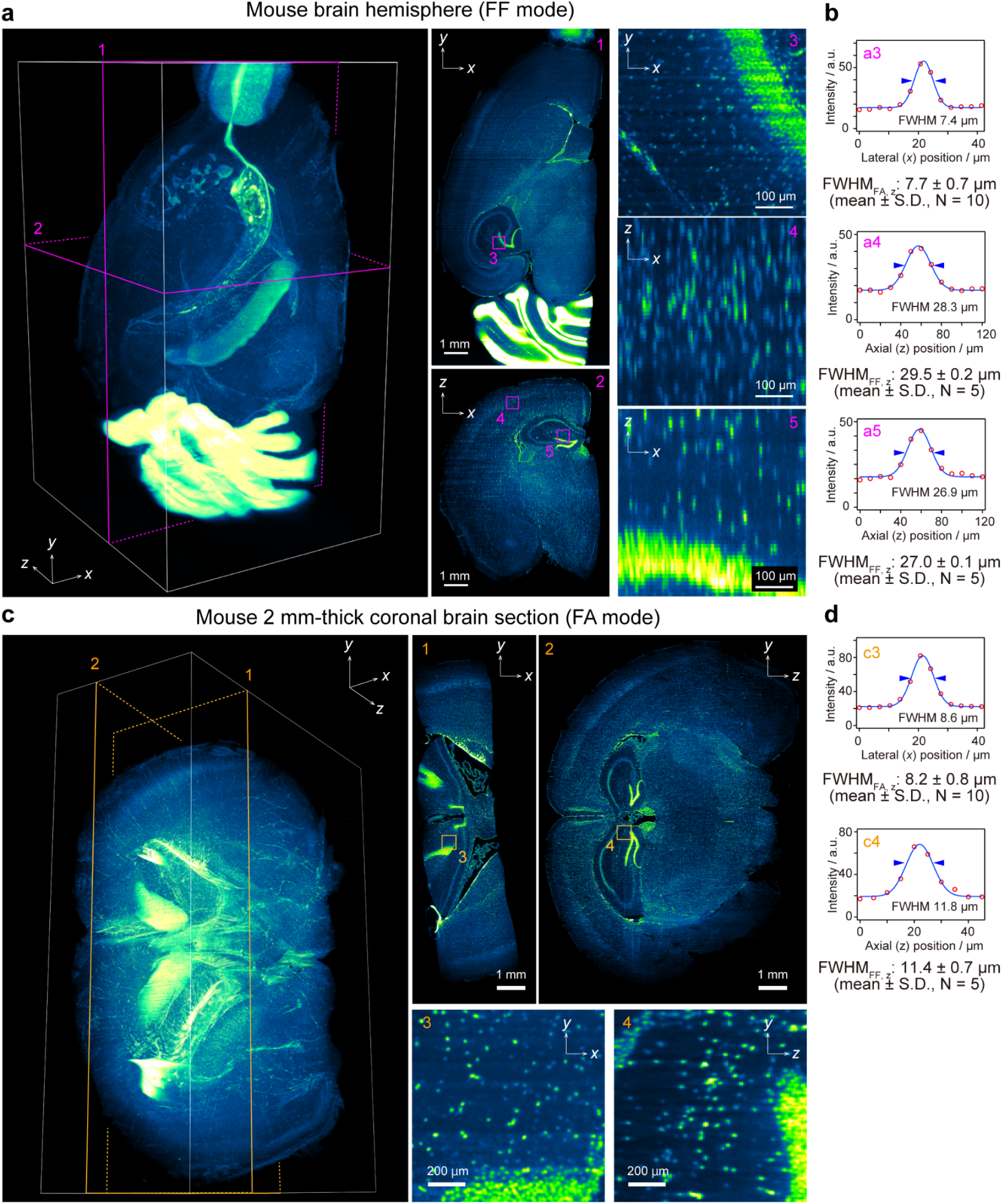
Optical specifications and imaging results. **a**. FF-mode imaging of a PI-stained mouse brain hemisphere. Voxel size: 3.45 × 3.45 × 10 μm^3^. Imaging speed: 180 sec. per 900 slices. Data size: 8.02 GB per 8-bit image stack. FFC was applied. The xy images illustrate a homogeneous image contrast across the FOV. The *yz* images indicate the elongation of PI-stained cell nuclei across the *z* direction due to the lower axial resolution. Note that the basic system did not include a device for eliminating the stripe shadow artifact (e.g., a galvo mirror) for simplification. Despite these technical limitations, the reconstituted 3D image is qualitatively sufficient for evaluating the whole sample structure. **b**. Lateral and axial spatial resolutions of PI-stained nuclei obtained with FF mode, estimated by the FWHM values of their intensity profiles. The axial FWHM values were consistent with the value measured by beads (**Fig. 1f**) and about quadrupled by the lateral FWHM of the PI signals. **c**. FA mode imaging of a PI-stained, 2 mm-thick mouse brain section. Voxel size: 3.45 × 3.45 × 5 μm^3^. Imaging speed: 372 sec. per 1860 slices per stack. Data size: 16.54 GB per 8-bit image stack. A total of four stacks with different light-sheet focus positions were collected, then TLS was applied. The image intensity was corrected by FFC. **d**. Lateral and axial spatial resolutions of PI-stained nuclei obtained by FA mode, estimated as in **b**. The *yz* images indicate that the FA mode provides a quality of image close to the isotropic resolution for the object with cell nuclei size across the *z* direction.

In conclusion, the basic configurations and specifications of descSPIM introduced here allows end-users to utilize a personalized LSFM, next to the bench for preparing cleared tissue samples. Furthermore, the descSPIM-basic is highly extendable and customizable via add-ons, such as a galvo for removing stripe-shape artifacts and near-infrared illumination-detection paths for live imaging. We also highlight its educational value: the users’ experience in constructing the basic equipment can aid in knowledge acquisition and alleviate the psychological burdens associated with advanced optics. Inspired by prior works such as openSPIM(*9*), mesoSPIM(*8*), and legolish/lemolish (https://lemolish.mystrikingly.com/), and adopting the spirit of open microscopy(*16*), our github website (https://github.com/dbsb-juntendo/descSPIM) provides all the instructions and the opportunity of community building. Almost 30 research groups have begun deploying the device internationally, indicating the solution meeting the unmet demand for broader tissue clearing applications.

## Acknowledgements

We thank the lab members at DBSB Juntendo, particularly Y. Wada for setting data analysis environments, S. Yanagi and D. Inahara for helping 3D CAD preparation. We also thank the Biomedical Research Core Facilities, Juntendo University Graduate School of Medicine, for technical assistance. This study was supported by PRIME (AMED to E.A.S., grant number JP20gm6210027); Project for Cancer Research and Therapeutic Evolution (P-CREATE) (AMED to E.A.S, grant number JP22ama221517); Research on Development of New Drugs (AMED to E.A.S., grant number JP21ak0101181); Brain/MINDS (AMED to E.A.S., grant number JP21wm0425003); CREST (JST to K.O, grant number JPMJCR20E4); JSPS KAKENHI grant-in-aid for scientific research (B) (to K.O. and E.A.S., grant number 22H02756 and 22H02824) and Grant-in-Aid for Transformative Research Areas - Platforms for Advanced Technologies and Research Resources “Advanced Bioimaging Support” (to E.A.S., grant number JP22H04926); Operating Costs Subsidies for Private Universities (to E.A.S.); Grants-in-Aid from UTEC-UTokyo, the Takeda Science Foundation, Nakatani foundation for advancement of measuring technologies in biomedical engineering, and Mochida Memorial Foundation for Medical and Pharmaceutical Research (to E.A.S.).

## Contributions

E.A.S. and K.O. designed the project and constructed the microscope. Y.S., and E.A.S. prepared the cleared and stained samples. K.O. acquired imaging data. T.O. and E.A.S. wrote the codes for image processing. Y.N. prepared the descSPIM Github website. K.O., T.O., and E.A.S. wrote the manuscript with input from all co-authors.

## Methods

### Tissue samples

We used 8-week-old male C57BL/6N mice (Japan SLC, Inc., Shizuoka, Japan). For sampling the tissues, the mice were sacrificed under the deep anesthetization with intraperitoneal administration of an overdose (> 100 mg/kg) of pentobarbital (pentobarbital sodium salt, nacalai tesque, Kyoto, Japan, #02095-04) and then transcardially perfused with PBS (occasionally supplied with ∽10 U/ml of heparin) and 4% paraformaldehyde in PBS. Then, the tissue samples were excised and postfixed with the same fixative for 8–24 h at 4 °C. The specimens were used for the subsequent clearing procedure or stocked at 4 °C in PBS with NaN_3_ until required for use. All experimental procedures and housing conditions of the animals were approved by the Animal Care and Use Committees of Juntendo University (1569-2022279 and 1372-2022211). All of the animals were cared for and treated humanely in accordance with the Institutional Guidelines and with the recommendations of the United States National Institutes of Health for experiments using animals.

### Tissue clearing and nuclear staining

CUBIC tissue clearing was performed according to our previous protocols(*13, 17*) with some optimizations for each sample. We used commercialized CUBIC-L and CUBIC-R+(N) reagents (TCI, #T3740 and #T3983). For PI staining, CUBIC-L-treated specimens were immersed in HEPES-NaCl buffer (10 mM HEPES: TCI, #H0396; 500 mM NaCl: TCI, #S0572; 0.05% NaN_3_: nacalai tesque, #31208-82) with 2-3 μg/mL Propidium iodide (PI) (DOJINDO, Kumamoto, Japan, #343-07461) at 37°C for 3-7 days according to the sample size.

### Microscopy construction

All parts list and the detailed information on construction and usage of descSPIM are opened in our Github website (https://github.com/dbsb-juntendo/descSPIM).

In brief, four-color laser lights (Cobolt, Stockholm, Sweden, Skyra, 488 nm/50 mW, 515 nm/50 mW, 561 nm/50 mW, and 647 nm/50 mW) were coupled into a single-mode fiber (Schäfter+Kirchhoff, Hamberg, Germany, PM series, effective NA = 0.072) and introduced to the excitation optical path. An achromatic convex lens (Thorlabs, Newton, NJ, C254-100-A, f = 100 mm) and two selectable cylindrical lenses (Thorlabs, LJ1144RM-A, f = 500 mm plano convex for the full-FOV mode; Thorlabs, ACY254-150-A, f = 150 mm achromatic for the fine-axial mode) were used for collimation and light-sheet formation, respectively. A CMOS camera (Thorlabs, S895MU, 7.45 × 14.1 mm^2^ imaging area, 3.45 × 3.45 μm^2^ pixel size, and 8.9 Megapixels) with a tube lens (Thorlabs, TTL100-A, f = 100 mm) and an 2X objective lens (Tholabs, TL2X-SAP, NA = 0.1, WD = 56.3 mm) were used for the detection optical path. One of Φ25 mm bandpass filters (Thorlabs, FBH520-40, 500-540 nm, FBH550-40, 530-570 nm, FBH600-40, 580 - 620 nm, or FBH700-40, 680 - 720 nm) or a longpass filter (Thorlabs, FBLH0550, 550 nm -) was placed in the detection path between the objective and the tube lenses. Motorized translation stages (Thorlabs, XR25P/M and Z825B) were adopted under the detection path and the sample stage. The sample was immersed in the applied clearing reagent or embedded in agarose(*13*) (http://cubic.riken.jp/) in a four-sided transmission cuvette (BK-7 glass cuvettes: GL Sciences, Tokyo, Japan, F10-G-10, 10 × 10 mm^2^; F10-G-20, 20 × 10 mm^2^, a polystyrene cuvette: BIO-RAD, Hercules, CA, #1702415, 10 × 10 mm^2^), and was placed on a suspended sample stage combined with a *θ* stage (Thorlabs, PR01/M). No oil chamber was used. *X* and *y* positions were manually adjusted by manual translation stages (Thorlabs, XR25P/M for fine *x* positioning, 1519/M and MVS05/M for rough and fine *y* positioning, respectively) before imaging.

### General imaging procedure

descSPIM imaging was performed with one-sided illumination. Eight-bit images were collected by scanning the sample in the z-direction as a movie file, the method inspired by MOVIE scan(*13*). Even though the camera’s maximal bit depth is 12 bits, we recommend eight-bit data collection to prevent I/O delays during movie imaging on a PC with midrange specifications. During the imaging procedure, the focus was maintained by two motorized stages that traced synchronously (**Supplementary Fig.2**). The synchronous speed correction value (the relative velocity of actuators) was theoretically calculated as follows: The focal length of the objective lens (*f*) can be expressed as a linear combination of the ratio of physical distance (*D*) to the refractive index of the optical path (*n*).

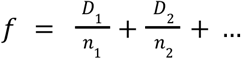

In the initial state of descSPIM, *f* can be described as;

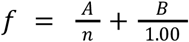

where *A* and *B* are the distances described in **Supplementary Fig.2**. When the sample position and the objective lens (*z*_sample_ and *z*_detect_) are moved in opposite directions (*dz*_sample_ and *dz*_detect_), *f* can be described as:

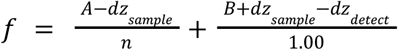

Therefore, *dz*_detect_ can be expressed as a function of *dz*_sample_, where

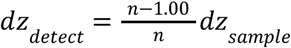

If CUBIC-R (*n* = 1.52) is utilized for clearing;

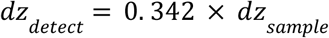

Therefore, the synchronous speed correction value is 0.342 in this case. This means that if the sample stage is moving at 100 μm/sec, the detection optics should be moving at 34.2 μm/sec. In the practical situation, however, the synchronous speed correction value can be influenced by the actuators’ capacity for weight and the PC’s specifications, resulting in a deviation from the theoretical estimate. We determined the practical correction value using the method illustrated in **Supplementary Fig. 4**.

When obtaining a z-stack image of a mouse hemisphere with FF mode (**Fig. 2a**), the velocity of the sample stage (*v*_stage_) was set to 50 μm/sec and the velocity of the detection optics (*v*_detect_) to 17.3 μm/sec on the actuator-associated software KINESIS™ (Thorlabs). The traveling distance of the sample stage was 10 mm (*dz*_stage_ = 10 mm, *dz*_sample_ = 3.46 mm). The time-lapse image was obtained with the exposure time of 200 ms. The resulting *xyt* image was converted to *xyz* format with a z-range of 10 mm and a *z*-interval (= *v*_stage_× exposure time) of 10 μm using ImageJ/Fiji (http://fiji.sc/Fiji). Due to the specification of the camera-associated software ThorCam™ (Thorlabs), files were split when they exceed 1 GB. Therefore, a final stack was generated using ImageJ/Fiji’s concatenate function. As for a mouse coronal section in FA mode (**Fig. 2b**), *v*_stage_ was set to 25 μm/sec and *v*_detect_ to 8.5 μm/sec. The traveling distance of the sample stage was 10 mm (*dz*_stage_ = 10 mm, *dz*_sample_ = 3.40 mm). The time-lapse images of multiple stacks for TLS were obtained with the exposure time of 200 ms. The resulting *xyz* images had a z-range of 10 mm and a z-interval of 5 μm. To visualize the fluorescent signals from PI, the 515 nm excitation laser light (3 mW output) and the 550 nm longpass filter were selected for measurements.

### PSF evaluation

Fluorescently-labeled beads (Nile Red, 1 μm diameter; Thermo Fisher Scientific) were diluted in the CUBIC-R+(N) agarose gel (1:1000, v/v) and filled into a cuvette for measurements. The fluorescent images of the Φ1 μm beads had a pixel size of 3.45 μm in both FF and FA mode. We acquired 300-μm-thick *z*-stacks at 10 μm intervals in FF mode and 120-μm-thick *z*-stacks at 3 μm intervals in FA mode, and used them to reconstruct *xyz* images of the beads. To visualize the fluorescent signals from the beads, the 515 nm excitation laser light (5 mW output) and the 550 nm longpass filter were selected for measurements. The FWHM values were evaluated by fitting their fluorescence intensity profiles around the central peak intensity using Gaussian functions. To avoid evaluation of intensity profiles derived from multiple fluorescent beads or position-dependent PSF elongations, we selected 10 of the dozens of FWHM values measured in a certain region of interest from the smallest values and evaluated their means and standard deviations.

### Adoption of the flat-field correction

The flat-field correction was incorporated from published literature(*14*) and implemented as an ImageJ macro. Briefly, 1 μM fluorescein was added to CUBIC-R reagent in a cuvette, and an image (reference image) was obtained using the same illumination mode (FF or FA) as the sample imaging except for exposure time of 2 s and bit-depth of 12-bit. Then, the median values in the *y* direction of the reference image were obtained and utilized to calculate the coefficient values for normalizing each median value to the peak median value in the *y* direction. The coefficient values were then applied to the sample stack in order to equalize the intensities along the *y*-axis.

### Adoption of the tiling light-sheet method

Tiling light-sheet imaging methods (TLS) with FA mode (**Fig. 2c**) was adopted from published literature(*15*) so that the eFOV (2× Rayleigh length) of the sheet illumination can cover the wider area along the x-axis after tiling. With a manual linear translation stage (Thorlabs, CTA1/M) affixed to the FA mode cylindrical lens (f = 150 mm), the light-sheet focus positions were moved along the x-axis by 500 μm in physical length (approximately 760 μm in CUBIC-R, RI = 1.52). In order to acquire multiple stacks at various light-sheet focus positions, the initial focal position was aligned at the center (or edge) of the sample. After collecting the first z-stack, the stage was moved 500 μm away and a second z-stack was collected, yielding z-stacks with focal points displaced by distance × refractive index value (less than twice the Rayleigh length in FA mode). These measurements were repeated to derive z-stacks for all of the sample’s regions of interest.

Tiling position in the x axis was identified using a custom ImageJ macro executing the process as follows: 1) A part of resliced x-z images from z-stacks that had different sheet focus positions were prepared. 2) The “Bandpass Filter” function of ImageJ was applied to a 50 × 100 pixel kernel. 3) A standard deviation (SD) was calculated for the resultant kernel area image. If the area contains axially focused high-contrast signals, the SD increases. 4) The SDs of the same kernel position from the two z-stacks were compared. 5) The point in x where the two SDs were inverted is identified as the tiling x-coordinate. Finally, the tiling was completed using the “combine” function of ImageJ at the calculated x-coordinate or using the fusion function of BigStitcher(*18*) with a 100-pixel overlap.

### General image processing for visualization

Acquired images were processed, reconstituted and visualized as a pseudo-colored image by using Fiji and NIS Elements (Nikon, Tokyo, Japan, version 5.02). The intensity was adjusted with the brightness and contrast functions. Occasionally, some images were cropped, and their orientations were adjusted to prepare figure panels.

**Supplementary Table 1.**
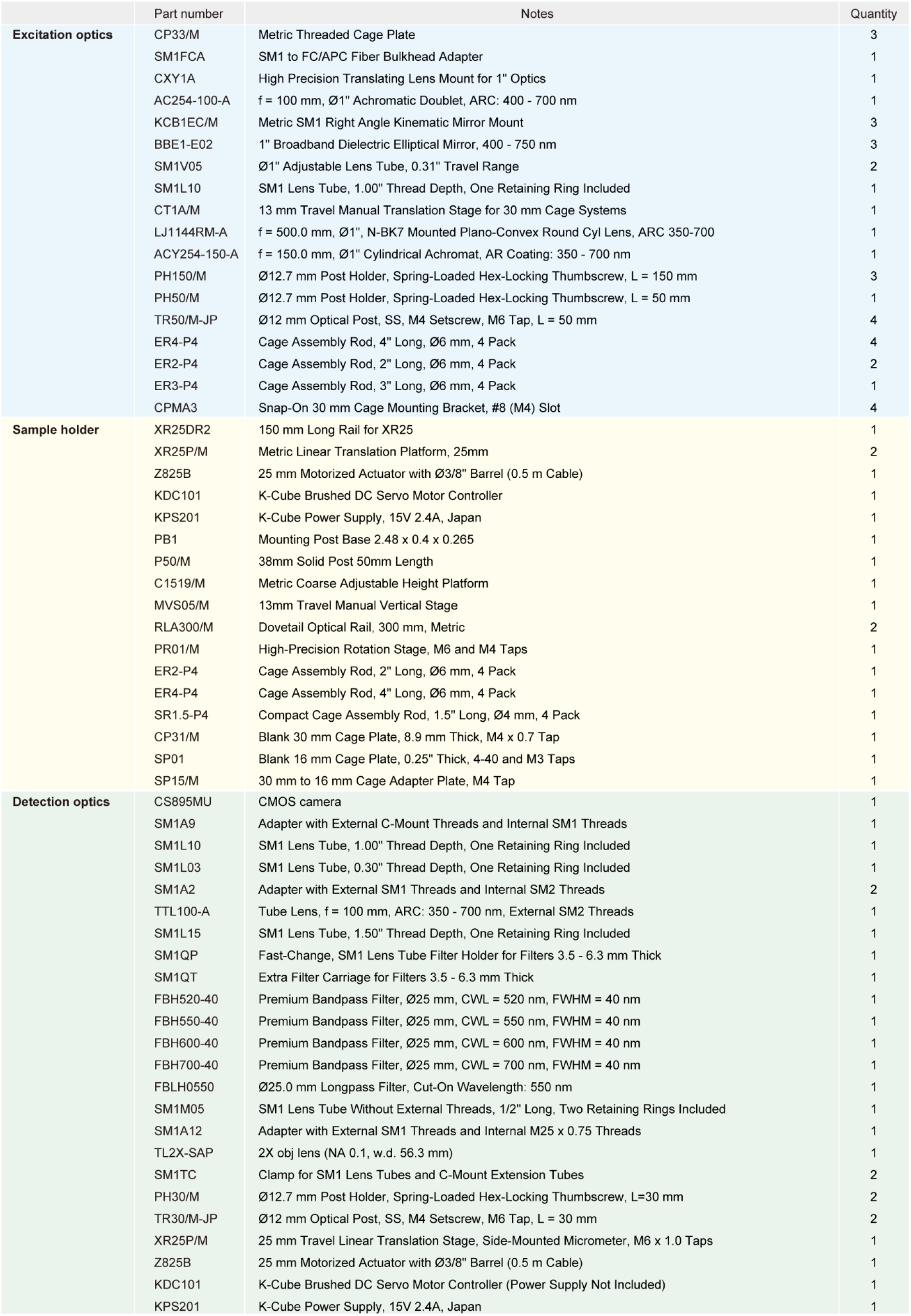
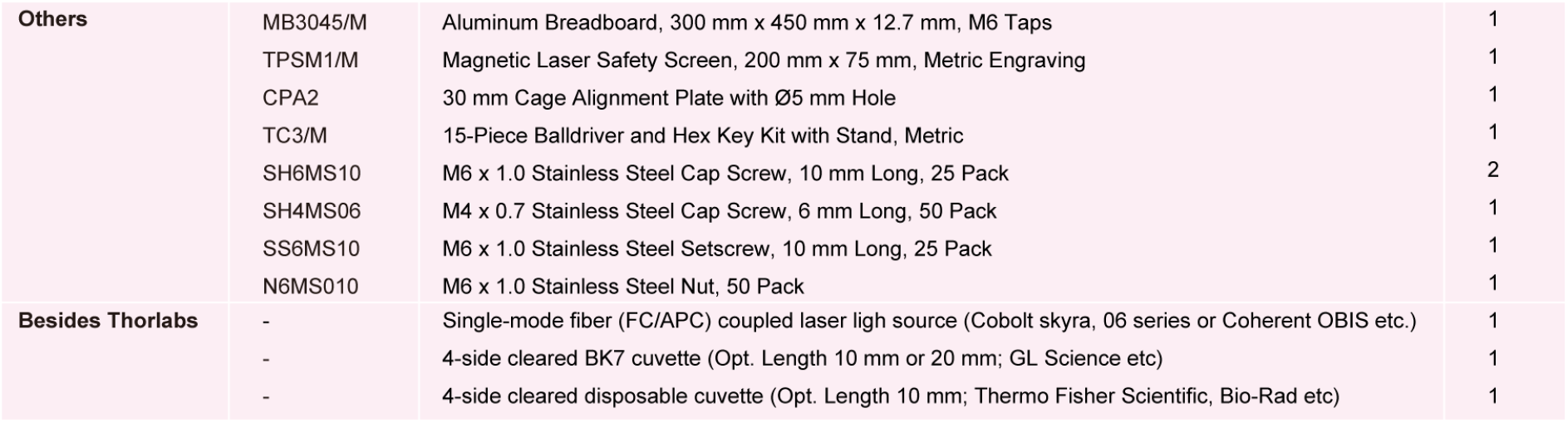
System components list of descSPIM.

|  | Part number | Notes | Quantity |
| --- | --- | --- | --- |
| Excitation optics | CP33/M | Metric Threaded Cage Plate | 3 |
|  | SM1FCA | SM1 to FC/APC Fiber Bulkhead Adapter | 1 |
|  | CXY1A | High Precision Translating Lens Mount for 1" Optics | 1 |
|  | AC254-100-A | f = 100 mm, Ø1" Achromatic Doublet, ARC: 400 - 700 nm | 1 |
|  | KCB1EC/M | Metric SM1 Right Angle Kinematic Mirror Mount | 3 |
|  | BBE1-E02 | 1" Broadband Dielectric Elliptical Mirror, 400 - 750 nm | 3 |
|  | SM1V05 | Ø1" Adjustable Lens Tube, 0.31" Travel Range | 2 |
|  | SM1L10 | SM1 Lens Tube, 1.00" Thread Depth, One Retaining Ring Included | 1 |
|  | CT1A/M | 13 mm Travel Manual Translation Stage for 30 mm Cage Systems | 1 |
|  | LJ1144RM-A | f = 500.0 mm, Ø1", N-BK7 Mounted Plano-Convex Round Cyl Lens, ARC 350-700 | 1 |
|  | ACY254-150-A | f = 150.0 mm, Ø1" Cylindrical Achromat, AR Coating: 350 - 700 nm | 1 |
|  | PH150/M | Ø12.7 mm Post Holder, Spring-Loaded Hex-Locking Thumbscrew, L = 150 mm | 3 |
|  | PH50/M | Ø12.7 mm Post Holder, Spring-Loaded Hex-Locking Thumbscrew, L = 50 mm | 1 |
|  | TR50/M-JP | Ø12 mm Optical Post, SS, M4 Setscrew, M6 Tap, L = 50 mm | 4 |
|  | ER4-P4 | Cage Assembly Rod, 4" Long, Ø6 mm, 4 Pack | 4 |
|  | ER2-P4 | Cage Assembly Rod, 2" Long, Ø6 mm, 4 Pack | 2 |
|  | ER3-P4 | Cage Assembly Rod, 3" Long, Ø6 mm, 4 Pack | 1 |
|  | CPMA3 | Snap-On 30 mm Cage Mounting Bracket, #8 (M4) Slot | 4 |
| Sample holder | XR25DR2 | 150 mm Long Rail for XR25 | 1 |
|  | XR25P/M | Metric Linear Translation Platform, 25mm | 2 |
|  | Z825B | 25 mm Motorized Actuator with Ø3/8" Barrel (0.5 m Cable) | 1 |
|  | KDC101 | K-Cube Brushed DC Servo Motor Controller | 1 |
|  | KPS201 | K-Cube Power Supply, 15V 2.4A, Japan | 1 |
|  | PB1 | Mounting Post Base 2.48 x 0.4 x 0.265 | 1 |
|  | P50/M | 38mm Solid Post 50mm Length | 1 |
|  | C1519/M | Metric Coarse Adjustable Height Platform | 1 |
|  | MVS05/M | 13mm Travel Manual Vertical Stage | 1 |
|  | RLA300/M | Dovetail Optical Rail, 300 mm, Metric | 2 |
|  | PR01/M | High-Precision Rotation Stage, M6 and M4 Taps | 1 |
|  | ER2-P4 | Cage Assembly Rod, 2" Long, Ø6 mm, 4 Pack | 1 |
|  | ER4-P4 | Cage Assembly Rod, 4" Long, Ø6 mm, 4 Pack | 1 |
|  | SR1.5-P4 | Compact Cage Assembly Rod, 1.5" Long, Ø4 mm, 4 Pack | 1 |
|  | CP31/M | Blank 30 mm Cage Plate, 8.9 mm Thick, M4 x 0.7 Tap | 1 |
|  | SP01 | Blank 16 mm Cage Plate, 0.25" Thick, 4-40 and M3 Taps | 1 |
|  | SP15/M | 30 mm to 16 mm Cage Adapter Plate, M4 Tap | 1 |
| Detection optics | CS895MU | CMOS camera | 1 |
|  | SM1A9 | Adapter with External C-Mount Threads and Internal SM1 Threads | 1 |
|  | SM1L10 | SM1 Lens Tube, 1.00" Thread Depth, One Retaining Ring Included | 1 |
|  | SM1L03 | SM1 Lens Tube, 0.30" Thread Depth, One Retaining Ring Included | 1 |
|  | SM1A2 | Adapter with External SM1 Threads and Internal SM2 Threads | 2 |
|  | TTL100-A | Tube Lens, f = 100 mm, ARC: 350 - 700 nm, External SM2 Threads | 1 |
|  | SM1L15 | SM1 Lens Tube, 1.50" Thread Depth, One Retaining Ring Included | 1 |
|  | SM1QP | Fast-Change, SM1 Lens Tube Filter Holder for Filters 3.5 - 6.3 mm Thick | 1 |
|  | SM1QT | Extra Filter Carriage for Filters 3.5 - 6.3 mm Thick | 1 |
|  | FBH520-40 | Premium Bandpass Filter, Ø25 mm, CWL = 520 nm, FWHM = 40 nm | 1 |
|  | FBH550-40 | Premium Bandpass Filter, Ø25 mm, CWL = 550 nm, FWHM = 40 nm | 1 |
|  | FBH600-40 | Premium Bandpass Filter, Ø25 mm, CWL = 600 nm, FWHM = 40 nm | 1 |
|  | FBH700-40 | Premium Bandpass Filter, Ø25 mm, CWL = 700 nm, FWHM = 40 nm | 1 |
|  | FBLH0550 | Ø25.0 mm Longpass Filter, Cut-On Wavelength: 550 nm | 1 |
|  | SM1M05 | SM1 Lens Tube Without External Threads, 1/2" Long, Two Retaining Rings Included | 1 |
|  | SM1A12 | Adapter with External SM1 Threads and Internal M25 x 0.75 Threads | 1 |
|  | TL2X-SAP | 2X obj lens (NA 0.1, w.d. 56.3 mm) | 1 |
|  | SM1TC | Clamp for SM1 Lens Tubes and C-Mount Extension Tubes | 2 |
|  | PH30/M | Ø12.7 mm Post Holder, Spring-Loaded Hex-Locking Thumbscrew, L=30 mm | 2 |
|  | TR30/M-JP | Ø12 mm Optical Post, SS, M4 Setscrew, M6 Tap, L = 30 mm | 2 |
|  | XR25P/M | 25 mm Travel Linear Translation Stage, Side-Mounted Micrometer, M6 x 1.0 Taps | 1 |
|  | Z825B | 25 mm Motorized Actuator with Ø3/8" Barrel (0.5 m Cable) | 1 |
|  | KDC101 | K-Cube Brushed DC Servo Motor Controller (Power Supply Not Included) | 1 |
|  | KPS201 | K-Cube Power Supply, 15V 2.4A, Japan | 1 |
| Others | MB3045/M | Aluminum Breadboard, 300 mm x 450 mm x 12.7 mm, M6 Taps | 1 |
|  | TPSM1/M | Magnetic Laser Safety Screen, 200 mm x 75 mm, Metric Engraving | 1 |
|  | CPA2 | 30 mm Cage Alignment Plate with Ø5 mm Hole | 1 |
|  | TC3/M | 15-Piece Balldriver and Hex Key Kit with Stand, Metric | 1 |
|  | SH6MS10 | M6 x 1.0 Stainless Steel Cap Screw, 10 mm Long, 25 Pack | 2 |
|  | SH4MS06 | M4 x 0.7 Stainless Steel Cap Screw, 6 mm Long, 50 Pack | 1 |
|  | SS6MS10 | M6 x 1.0 Stainless Steel Setscrew, 10 mm Long, 25 Pack | 1 |
|  | N6MS010 | M6 x 1.0 Stainless Steel Nut, 50 Pack | 1 |
| Besides Thorlabs | - | Single-mode fiber (FC/APC) coupled laser light source (Cobolt skyra, 06 series or Coherent OBIS etc.) | 1 |
|  | - | 4-side cleared BK7 cuvette (Opt. Length 10 mm or 20 mm; GL Science etc) | 1 |
|  | - | 4-side cleared disposable cuvette (Opt. Length 10 mm; Thermo Fisher Scientific, Bio-Rad etc) | 1 |

**Supplementary Fig. 1.**
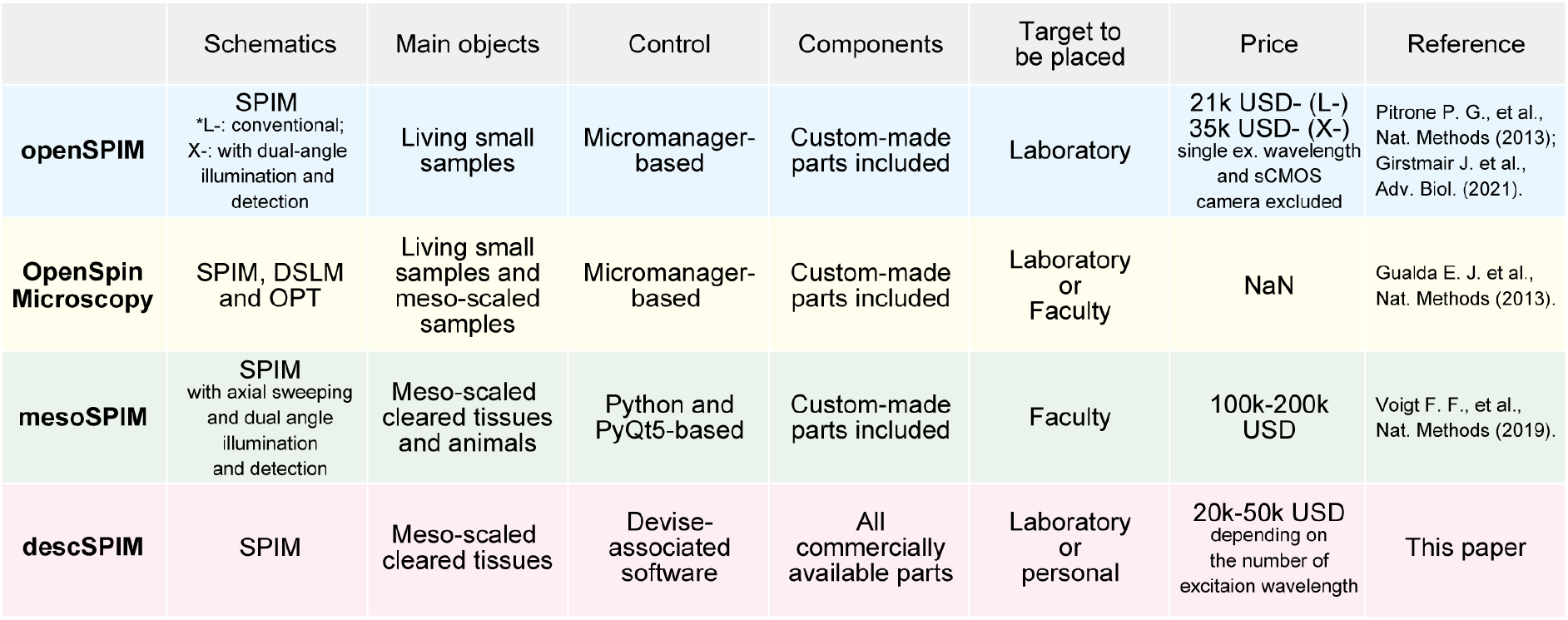
Design and comparison of descSPIM. A chart comparing open-source light-sheet microscopy. descSPIM is the only instrument capable of balancing cleared tissue imaging with an affordable price. descSPIM also excluded the use of custom-made components for user convenience.

**Supplementary Fig. 2.**
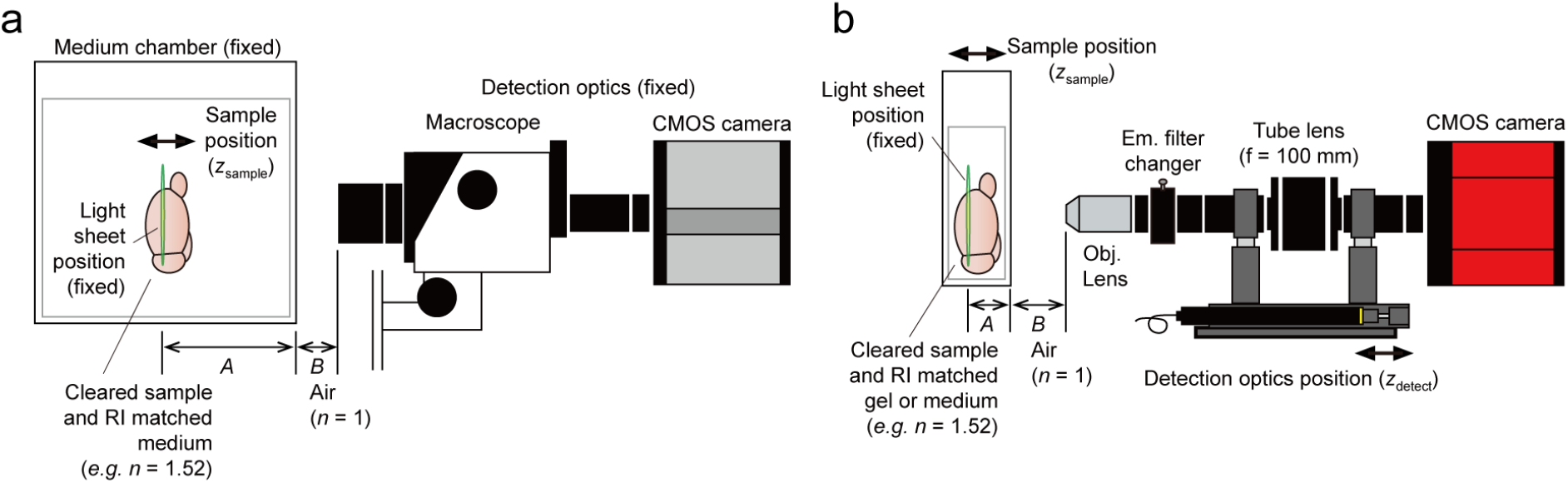
Implementation of focus tracing. **a**. A conventional system with a positionally fixed medium chamber fulfilled with a clearing reagent or a RI-matched immersion oil (e.g., *n* = 1.52). The sample is moved along the z-axis within the chamber. In this case, the ratio of *A* (the distance from the chamber wall to the light-sheet illumination, with RI of the immersion reagent) and *B* (the distance from the objective lens to the chamber wall, with RI of the air (1.0)) is fixed. **b**. descSPIM sample imaging method employing a cuvette as a sample container. In this case, due to the movement of the sample chamber (the cuvette) during imaging, the ratio of *A* and *B* is altered, resulting in defocus. descSPIM applies synchronized movement of the sample stage (z_sample_) and the detection optics (z_detect_) to prevent the defocus. See **Supplementary Fig. 4** and **Methods** for details on how to calculate the synchronous speed correction value (the relative velocity of two actuators).

**Supplementary Fig. 3.**
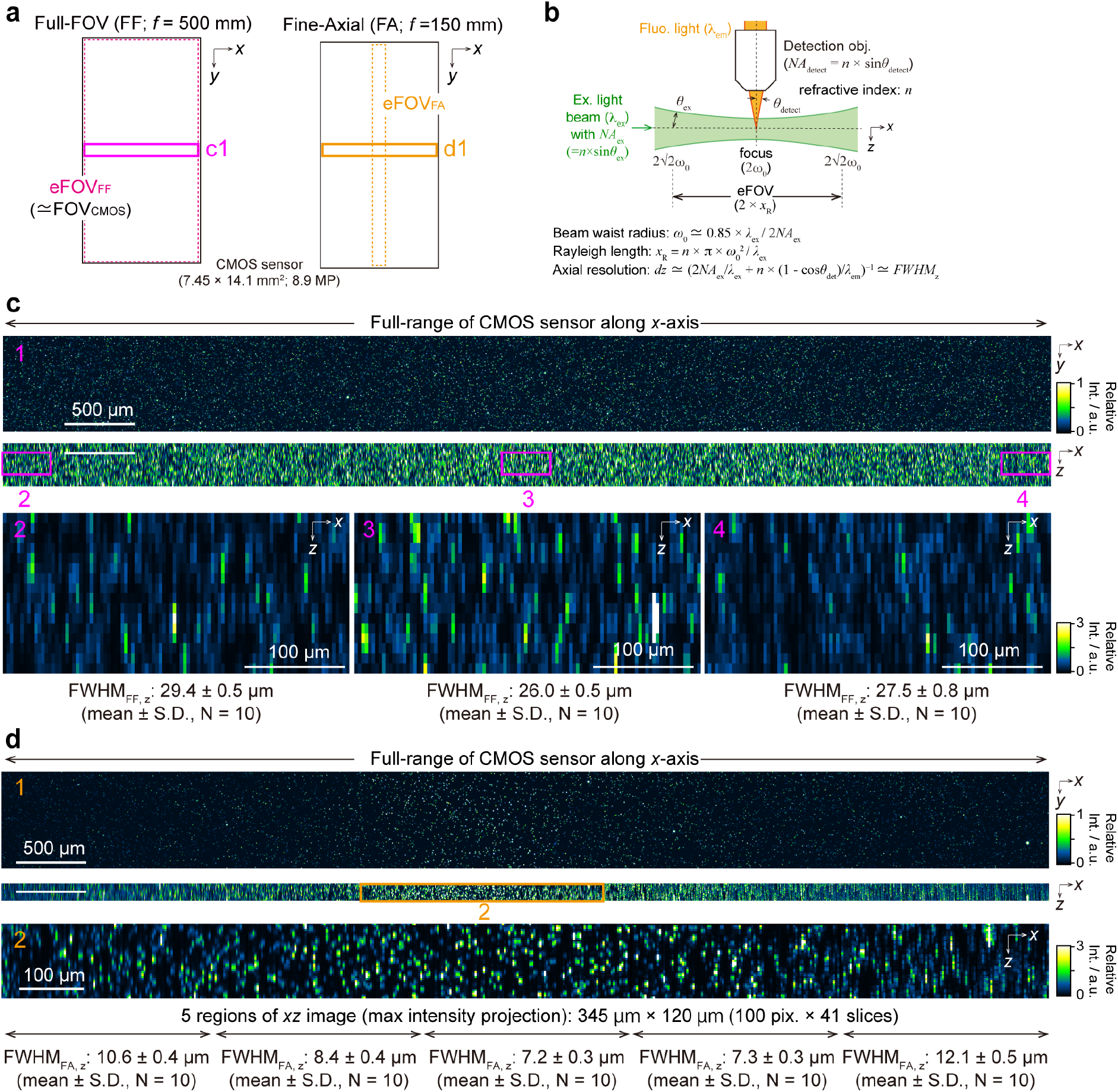
Detailed PSF calculation across the light-sheet illumination. **a**. The entire CMOS sensor area and assumed light-sheet coverages by FF and FA modes are shown. b1 and c1 indicate the places enlarged in **c** and **d**, respectively. eFOV: effective field of view. **b**. Definition of light-sheet illumination parameters from literature(*12*). The NA value of the cylindrical lens can be calculated with an axial resolution value (approximated as FWHM). eFOV, defined as 2 × Rayleigh length, can be then calculated with the NA value. **c**. Enlarged lateral (*x*-*y*) and axial (*x*-*z*) views of the microbeads (Φ1 μm) image (max intensity projection of 300 μm in *z* range and 690 μm in *y* range) across the full-range x-axis of the CMOS sensor, captured with FF mode. Axial FWHMs measured at the center and margins indicate that the light-sheet thickness is nearly homogeneous (approximately 26–29 μm) across the range. **d**. Enlarged lateral (*x*-*y*) and axial (*x*-*z*) views of the microbeads image (max intensity projection of 120 μm in *z* range and 690 μm in *y* range) captured with FA mode. At the sheet-focused position (the center of box 2), the axial FWHM is approximately 7.2 μm. The eFOV can be regarded up to the area where the FWHM is under ∽10.2 μm (√2 × 7.2 μm, based on the definition of Rayleigh length). The image indicates that the FWHM ranges between 7.2–10.2 μm over a width of 300 pixels. An estimate of the eFOV from the effective NA value is 830 μm, which is consistent with the result in panel 2.

**Supplementary Fig. 4.**
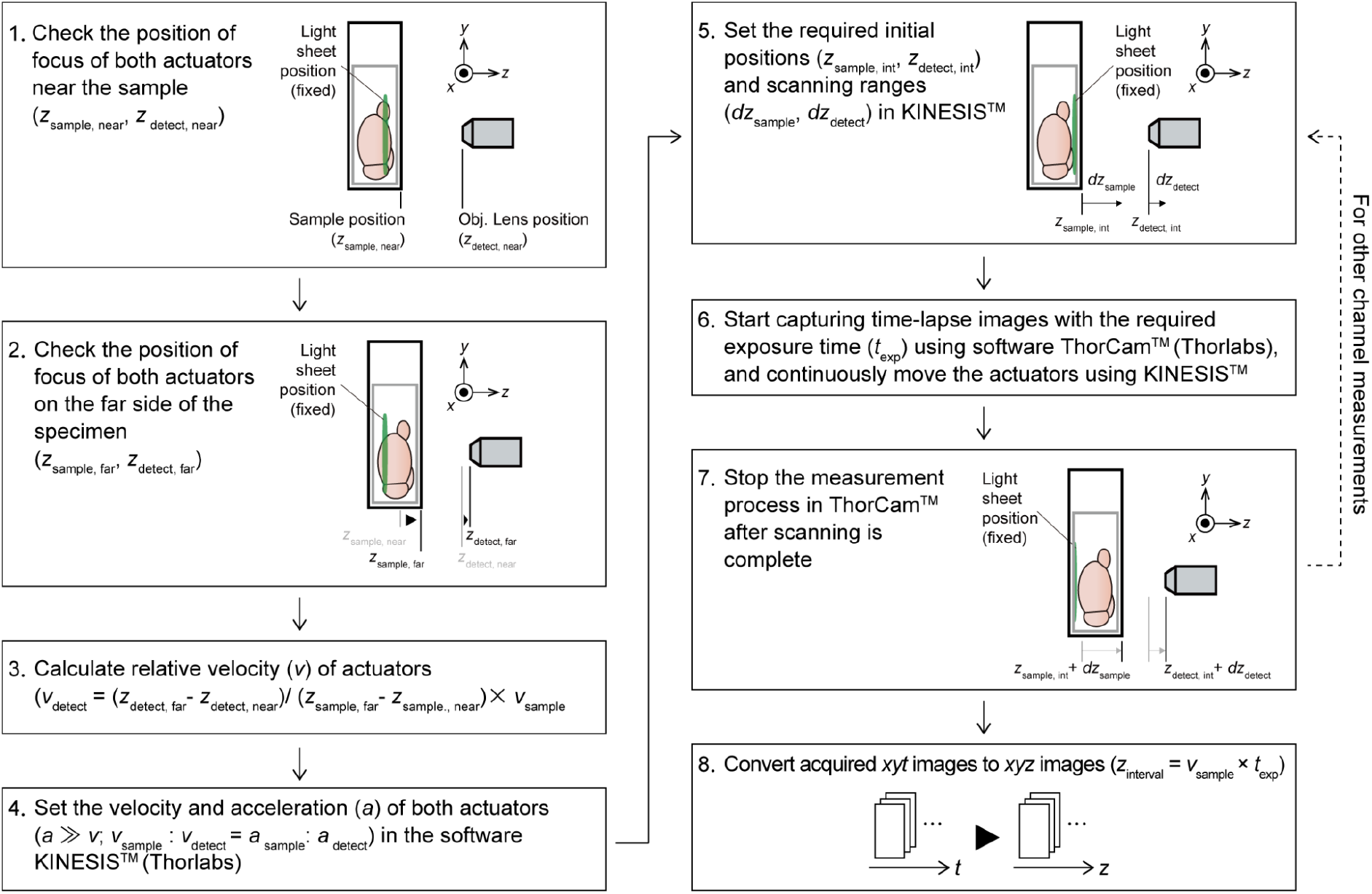
Operation workflow. The steps 1–3 are for calculating the synchronous speed correction value (relative velocity of actuators). Although this value should be theoretically estimated as in **Supplementary Fig. 2**, it can be influenced by the actuators’ capacity for weight and the PC’s specifications (see **Methods**). For calculating the value in the practical situation, an operator moves the sample stage to the z-stack start position and gets the stage position. Then, the operator moves the detection objective to the focused position, and also gets the actuator position (step 1). The operator repeats the same steps at the z-stack end position (step 2) and calculates the relative velocity of the actuators (step 3). The velocities, accelerations, initial positions of these two actuators, and their scanning ranges are set on the associated software (KINESIS, Thorlabs) (steps 4, 5). After the laser illumination is turned on, the operator starts image acquisition by clicking the start buttons of the KINESIS and the camera-associated software (ThorCam, Thorlabs). The z-stack is collected as a time-lapse (*xy-t*) file (steps 6, 7). Finally, the acquired movie is converted to a z-stack (*xy-z*) file with ImageJ/Fiji. The z-interval is calculated as the moving speed of the sample stage actuator (*v*_sample_) × exposure time (*t*_exp_) (step 8).

**Supplementary Fig. 5.**
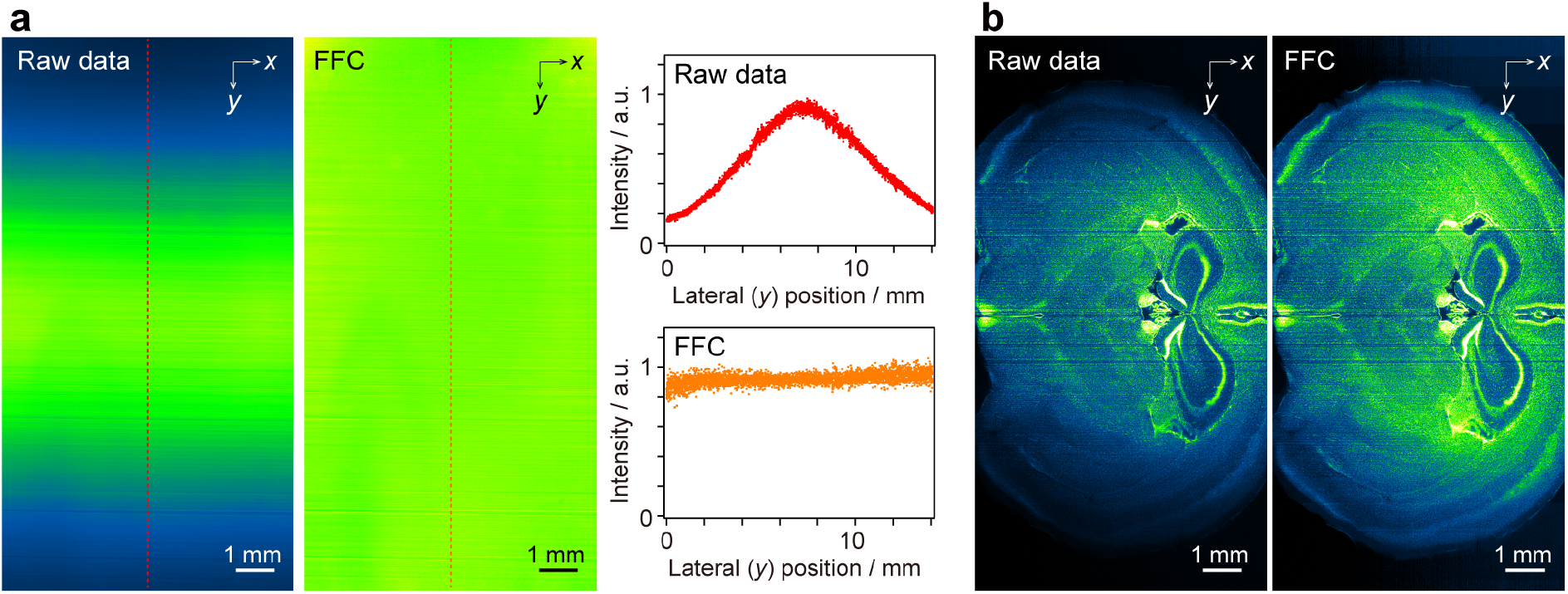
Flat-field correction (FFC) of Gaussian illumination. **a**. FFC implementation(*14*). A fluorescein-dissolved clearing reagent was imaged as a reference. The gaussian-shaped intensity profile was converted into a flat intensity profile by our custom imageJ macro code (see **Methods** and our Github website: https://github.com/dbsb-juntendo/descSPIM). **b**. An FFC example applied to the actual sample image. A coronal image of a 2 mm-thick mouse brain section stained with PI was obtained, and the FFC was applied.

**Supplementary Fig. 6.**
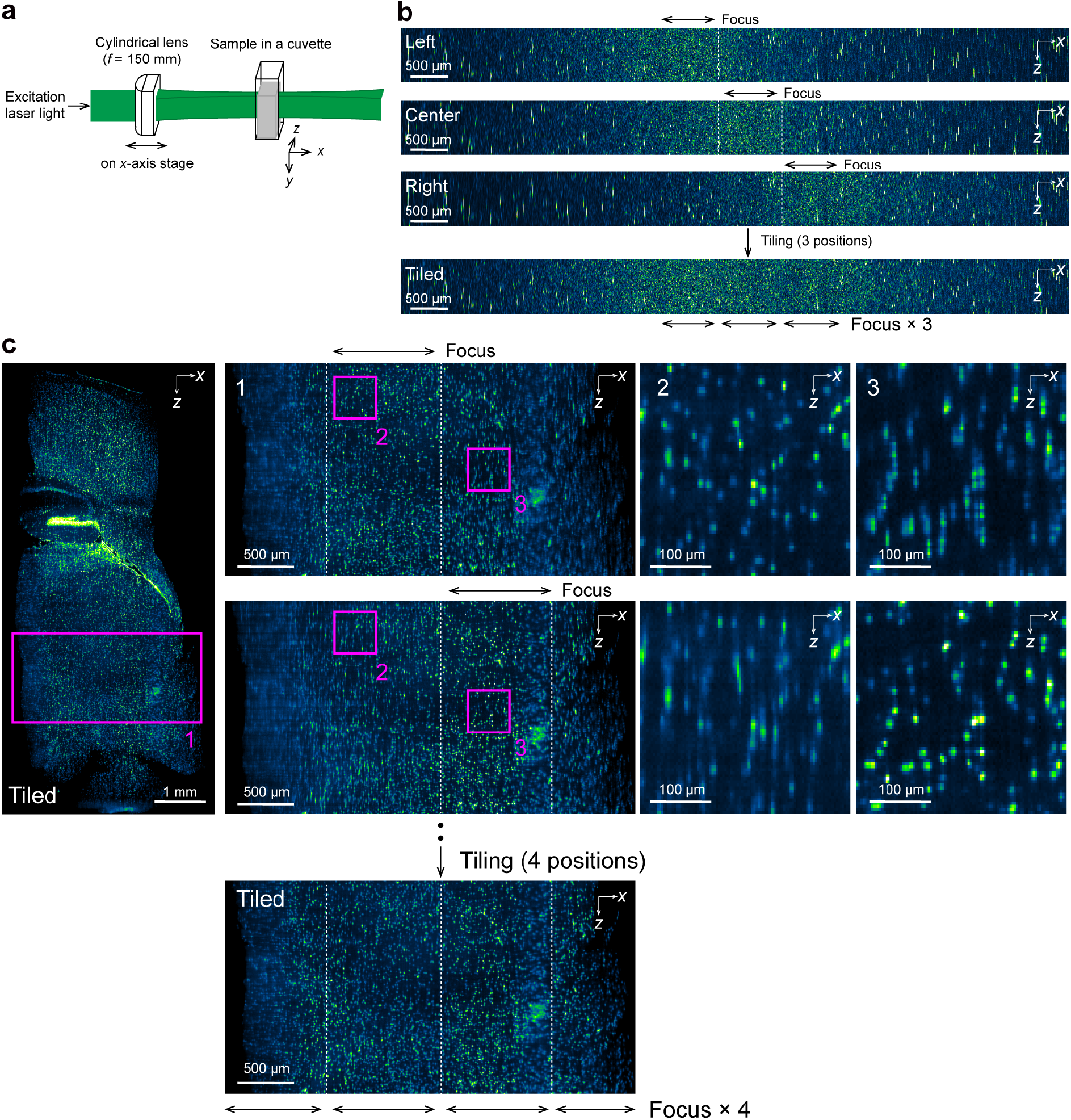
Adaptation of tiling light-sheet method. **a**. The imaging scheme. descSPIM adopts the tiling light-sheet method(*15*) by manually moving the cylindrical lens (f = 150 mm) for FA illumination mode. **b**. TLS procedure. The axial image of the bead signals (max intensity projection of 690 μm in *y* range) showed a high signal contrast area corresponding to the eFOV (practical 2× Rayleigh length) of the sheet illumination. Based on the estimation in **Supplementary Fig. 3**, we moved the cylindrical lens from the left (proximal side of the lens) to the right (distal side of the lens) by 500 μm in the air (corresponding to 760 μm in the medium with RI = 1.52) in each step. The tiling position was identified based on the signal contrast calculation with the Fourier function of ImageJ (see **Methods**). **c**. An example of TLS adoption for PI-stained 2 mm-thick mouse brain data. The dotted lines in **b** and **c** show the tiling positions on the x axis.

## Supplementary Movies

Supplemental movie 1. PI-stained mouse brain hemisphere, shown in **Fig. 2a**.

Supplemental movie 2. PI-stained 2-mm thick mouse brain section, shown in **Fig. 2c**.

